# Insects pollinated flowering plants for most of angiosperm evolutionary history

**DOI:** 10.1101/2023.01.31.526530

**Authors:** Ruby E. Stephens, Rachael V. Gallagher, Lily Dun, Will Cornwell, Hervé Sauquet

## Abstract

- Pollination is a fundamental process driving the speciation of angiosperms (flowering plants). Most contemporary angiosperms are insect pollinated, but abiotic pollination by wind or water and vertebrate pollination by birds or mammals occurs in many lineages. We model the evolution of pollination across angiosperms and quantify the timing and environmental associations of pollination shifts.
- We use a robust dated phylogeny and trait-independent species-level sampling across all families of angiosperms to model the evolution of pollination modes. Data on the pollination system or syndrome of 1160 species were collated from primary literature.
- Angiosperms were ancestrally insect pollinated, and insects have pollinated angiosperms for approximately 86% of angiosperm evolutionary history. Wind pollination evolved at least 42 times, with few reversals back to animal pollination. Transitions between insect and vertebrate pollination were more frequent: vertebrate pollination evolved at least 39 times from an insect pollinated ancestor with at least 26 reversals. The probability of wind pollination increases with habitat openness (measured by Leaf Area Index) and with distance from the equator.
- Our reconstruction of pollination across angiosperms sheds light on a key question in angiosperm macroevolution, highlighting the long history of interactions between insect pollinators and angiosperms still vital to global biodiversity today.

## 1 Introduction

Pollination is a fundamental ecological process that has influenced the diversification of many seed plant families throughout evolutionary history (Ollerton *et al*., 2019; Asar *et al*., 2022). Both gymnosperms and angiosperms depend on pollination to reproduce sexually, with pollen transfer effected by insects, vertebrates, wind or water as vectors (Faegri & van der Pijl, 1979). Shifts between different pollinators are often implicated in the speciation of closely related plants, and in the angiosperms pollination shifts have driven the evolution of the vast array of floral forms present today (van der Niet & Johnson, 2012; van der Niet *et al*., 2014).

Precisely how the first angiosperms were pollinated, and how pollination modes have evolved through time, remains a key question in angiosperm macroevolution (Sauquet & Magallón, 2018). The majority of angiosperms are pollinated by animals, especially insects (e.g. bees, flies, wasps, moths, butterflies, beetles, thrips) but also vertebrates (e.g. birds, bats, lizards, small mammals) (Faegri & van der Pijl, 1979; Ollerton *et al*., 2011). Indeed, although some flowers self-pollinate, up to a third of angiosperms set no seed at all without animal pollination (Rodger *et al*., 2021). However, abiotic pollination by wind or water also occurs in many diverse plant lineages, and wind pollination is estimated to have evolved at least 65 times across the angiosperms (Linder, 1998; Ackerman, 2000). Combined pollination by animals and wind (ambophily) is also found in many unrelated lineages, and may be more common than currently reported as it is rarely tested for in pollination studies (Culley *et al*., 2002; Abrahamczyk *et al*., 2022a).

It is widely believed that the most recent common ancestor of the angiosperms was insect pollinated (Hu *et al*., 2008; Labandeira & Currano, 2013; Gottsberger, 2016; Asar *et al*., 2022). This is supported by the predominance of insect pollination in extant early-diverging angiosperms and in fossil seed plants (Hu *et al*., 2008; Friis *et al*., 2011; Asar *et al*., 2022), though extant early-diverging angiosperms also include wind pollinated (e.g. *Trithuria submersa*, Taylor *et al*., 2010) and ambophilous taxa (e.g. *Amborella trichopoda*, Thien *et al*., 2003). Ancestral pollination is yet to be explored on the full angiosperm phylogeny, however, and questions persist about the timing and tempo of pollination mode evolution. For instance, it is not yet known when shifts to wind pollination occurred, and whether these were as common as shifts between insect and vertebrate pollinators. Yet to be explored also is the frequency of reversals from wind back to animal pollination, and how the ancestors of all major angiosperm clades may have been pollinated.

The environmental conditions that have accompanied shifts between pollination modes across angiosperm evolution also remain unclear. Despite macroecological evidence that wind pollination decreases towards the equator (Ollerton *et al*., 2011; Rech *et al*., 2016), evolutionary studies show no relationship between wind pollination and geographical distribution (Friedman & Barrett, 2008). Wind pollination appears to have evolved more often in open habitats, however, where pollen is more easily airborne, and is less common today in warm, wet and species rich environments (Friedman & Barrett, 2008; Rech *et al*., 2016). Given these mixed ecological and evolutionary relationships, whether shifts to wind pollination have consistently been associated with shifts between habitats or major biomes during angiosperm evolution bears further investigation.

Here we quantify major changes in the evolution of pollination modes across a robust dated phylogeny (Ramírez-Barahona *et al*., 2020), and an unprecedented suite of trait-independent species-level observations across all families and major subfamilies of angiosperms. We estimate the rate and timing of transitions between insect, vertebrate, wind and water pollination, and reconstruct the ancestral pollination modes of major angiosperm lineages. We also use this dataset to quantify macroecological patterns of animal versus wind pollination in a phylogenetic framework. Specifically, we ask whether emergent relationships between wind pollination and latitude or habitat openness (as measured by Leaf Area Index, LAI) remain when angiosperm evolutionary history is considered.

## 2 Materials and Methods

### 2.1 Scoring pollination mode

Pollination system or syndrome was scored for 1201 species across 434 plant families contained in the angiosperm phylogeny of Ramírez-Barahona *et al*. (2020). Where possible pollination was scored at species level (n = 1025), cross-checked against what is known of pollination in that genus and family, especially from the Kubitzki series (Kubitzki *et al*., 1993–2018). Where no information was available for a particular species they were scored at genus (n = 131), or family (n = 4) level. We obtained pollination data for 1160 of 1201 taxa, using the best available evidence to score the pollination system (n = 432) or syndrome (n = 728) for each taxon. Where possible, explicit studies of pollination ecology in a taxon’s native range were preferred (n = 239), especially if these involved explicit tests for the occurrence of wind pollination (45 of these). Where these data were not available, we used records of floral visitation in combination with an interpretation of species floral syndrome (n = 193).

Where no field observations had been recorded, we interpreted species floral syndrome sensu Faegri & van der Pijl (1979) (n = 728). Although pollination syndromes can be inaccurate at fine taxonomic levels (Ollerton *et al*., 2009; van der Niet, 2021), they are effective predictors of the broad pollination groups used here (Rosas-Guerrero *et al*., 2014; Dellinger, 2020), especially wind pollination which has a well-defined suite of traits (Friedman & Barrett, 2008). Floral syndrome was interpreted from species descriptions, illustrations and images from various sources, including iNaturalist and eFloras (eFloras, 2022; iNaturalist, 2022), and informed by pollination syndrome data from trait databases including TRY (Kattge *et al*., 2020), BiolFlor (Kühn *et al*., 2004), and AusTraits (Falster *et al*., 2021). Full references are available in Supporting Information.

Floral syndrome was scored by considering all available evidence. To separate wind from animal pollinated flowers we assessed traits in Table 1 of Friedman & Barrett (2009) and pollen as described in Hu *et al*. (2008), particularly perianth size and colour, gynoecium size and shape, pollen and floral rewards (Table 1). To separate insect from vertebrate pollination syndromes we considered floral size and the robustness of floral parts, nectar quantity and the accessibility of floral rewards to different pollinators (e.g. the presence of poricidal anthers which only release pollen when vibrated by bees in buzz-pollinated flowers (Pritchard & Vallejo-Marín, 2020), Table 1). Water pollination was considered in the rare cases where plants had an aquatic habit and flowered near or under water (Ackerman, 2000).

**Table 1.** Some of the key floral traits used to assign species pollination syndromes, in addition to all other available evidence.

| Trait | Wind | Insect | Vertebrate |
| --- | --- | --- | --- |
| Scent | Absent | Often present | Present (mammal) or absent (bird) |
| Nectar | Absent | Often present | Present, abundant |
| Pollen | Dry, smooth, small, abundant, easily airborne | Sticky, clumped, larger, sometimes available via poricidal anthers | Sticky, clumped, larger |
| Perianth | Plain or reduced | Often present and brightly coloured | Present, robust |
| Gynoecium | Feathery or with increased surface area | - | - |
| Sexuality | Often unisexual | Often bisexual | Often bisexual |

Flowers were scored as polymorphic where there was evidence for more than one pollination mode, or where they were pollinated by animals but it was unclear whether this was vertebrate or insect pollination (n = 76). Where there was no evidence that pollination via external vectors occurred (in clonal or autogamous species) or no information was available these species were left as missing data (n = 41). Our final dataset included pollination information for 1160 of the 1201 species in 433 families (all except Hoplestigmataceae) in the Ramírez-Barahona *et al*. (2020) angiosperm tree. Fully referenced data are available at https://doi.org/10.5281/zenodo.7592528.

### 2.2 Ancestral state reconstructions and stochastic character mapping

Data processing and analysis was completed in R version 4.1.3 (R Core Team, 2022) using packages including the *tidyverse* collection (Wickham *et al*., 2019), *ape* version 5.6.2 (Paradis & Schliep, 2019), *corHMM* version 2.8 (Boyko & Beaulieu, 2021) and *phytools* version 1.0.3 (Revell, 2012).

For analyses we used a dated phylogeny from Ramírez-Barahona *et al*. (2020), specifically the maximum clade credibility time-tree reconstructed in BEAST using the ‘relaxed calibration strategy’ with one prior constraint on the crown age of angiosperms and 238 fossil-based minimum age constraints (‘RC-complete analysis’). To reconstruct ancestral pollination modes and estimate rates of transition between pollination modes we compared two Markov models via a maximum likelihood approach in corHMM: Equal Rates (ER) where all transition rates are equal and All Rates Different (ARD) where all transition rates differ. Pollination modes were separated into four states: wind, water, insect and vertebrate. Ancestral state reconstructions were run first with a model distinguishing among these four states (4-state model) and then with various simpler models combining these states into fewer categories (e.g. abiotic (wind or water) versus animal (insect or vertebrate) pollination, Supporting Information Table S1) to reduce the number of parameters in the models and to allow the possibility of ancestral polymorphisms. corHMM analyses were run with 10 random restarts using the default Yang root prior (Yang, 2006) and marginal node state reconstruction.

To estimate the timing and number of transitions between different pollination modes we used stochastic character mapping via the makeSimmap function in *corHMM* (1000 simulations across the tree for each state combination considered). Stochastic character mapping assessed all pollination states (wind-water-insect-vertebrate), as well as wind versus animal and vertebrate versus insect pollination. Taxa missing data were dropped from the phylogeny for wind-animal and vertebrate-insect analyses. To assess the impact of dating uncertainty on these results, we also ran stochastic character mapping across 1000 trees sampled from the posterior of Ramírez-Barahona *et al*. (2020), with 100 simulations on each tree.

### 2.3 Environmental correlates of wind pollination

To test for relationships between spatial and environmental variables and wind pollination we sourced occurrence data (latitude-longitude coordinates) from the Global Biodiversity Information Facility (GBIF) for the 1201 taxa in our dataset (filtered from GBIF.org, 2022). Occurrences were cleaned by retaining only records with an associated herbarium specimen collected since 1980, and using the package CoordinateCleaner version 2.0-20 (Zizka *et al*., 2019) to remove geographic outliers, non-land records, country-coordinate mismatches, points near known herbaria, botanic gardens, and country capitals, duplicated records, and points with identical longitude and latitude. Cleaning used the computational resources from the Katana computing cluster at UNSW (https://doi.org/10.26190/669x-a286).

Using occurrence data, we calculated the absolute mean latitude of each species range (distance from the equator) and a metric of habitat openness based on Leaf Area Index (LAI) across the range. LAI is defined as one half of the total green leaf area per unit horizontal ground surface area and is measured at frequent intervals to track plant growth responses, which change with seasonal and climatic conditions (Fang *et al*., 2019). LAI is highest in densely forested areas such as the Amazon rainforest and lowest in arid open areas such as the Sahara desert, and is an objective continuous measure of the openness of species habitat (Liu *et al*., 2012; Fang *et al*., 2019). We took one average of LAI from GLOBMAP data for July 1981-December 2020 (Liu *et al*., 2012), and extracted mean LAI for each GBIF occurrence, then averaged these for each species to provide a metric of species mean habitat openness.

To test whether wind pollination is more frequently associated with higher latitudes or open habitats we used phylogenetic logistic regression fitted in *phylolm* version 2.6.2 (Tung Ho & Ané, 2014) following the Ives and Garland method (Ives & Garland, 2010). Uncertainty of parameter estimates was estimated by 100 parametric bootstrap samples. 180 taxa with missing or polymorphic data were excluded from this analysis, giving final sample size n = 1021.

To test whether the evolution of wind pollination in angiosperms was correlated with shifts between major biomes we ran Pagel’s (1994) models of correlated evolution, see Supporting Information Notes S1 for details.

## 3 Results

Angiosperms were reconstructed as ancestrally insect pollinated, with all models supporting insect or animal pollination as the most likely ancestral state at the root of the angiosperm tree (Figure 1, Supporting Information Table S1). Results hereon are from the wind, water, insect, vertebrate 4-state model, for which the ARD model had highest support (SI Table S1).

**Figure 1.**
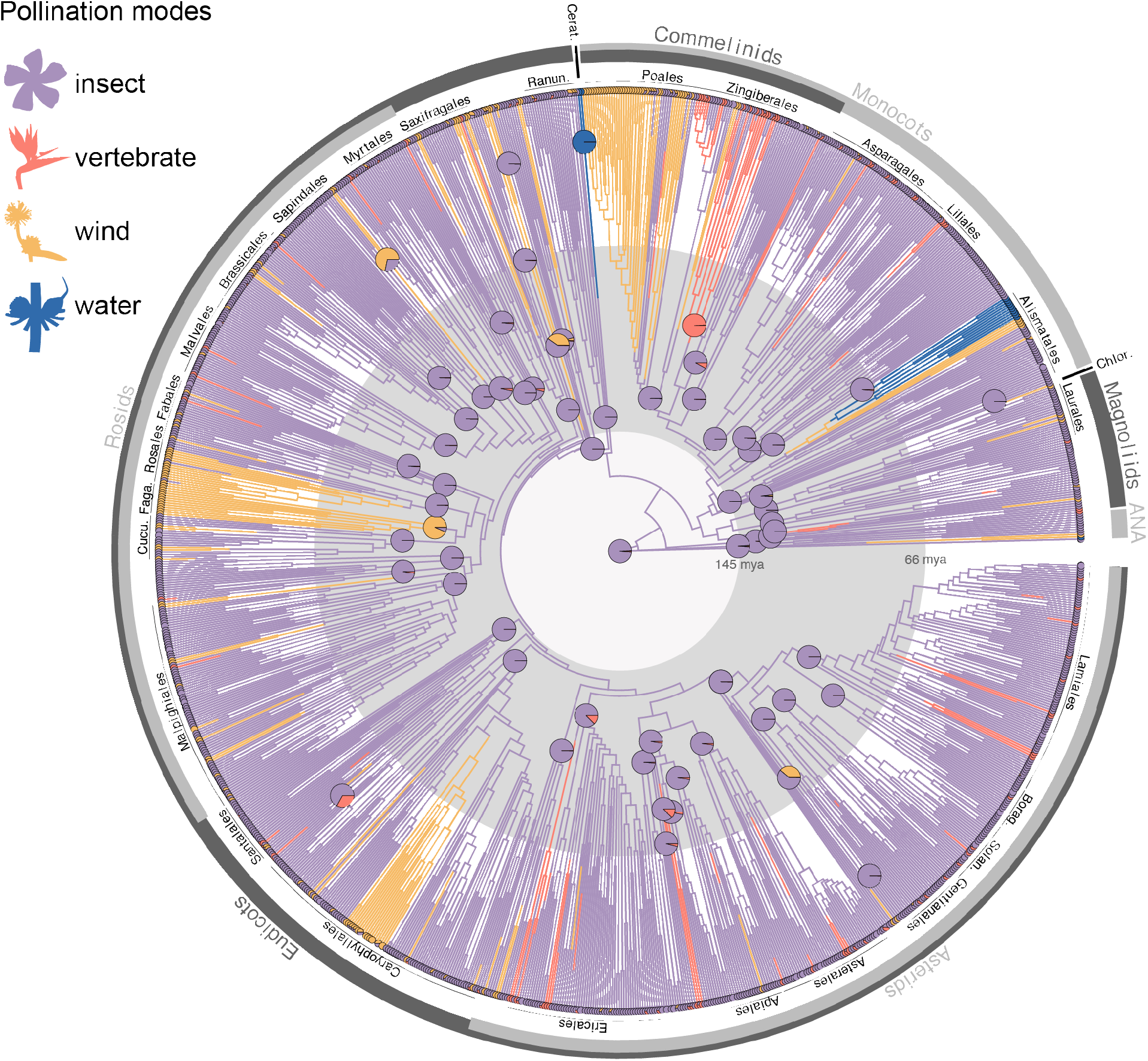
The macroevolution of pollination modes across angiosperms, showing the proportional marginal likelihood of pollination mode at the ancestral nodes for each angiosperm order (n = 64) from the 4-state ARD model. Colour along tree branches indicates pollination mode from one randomly selected stochastic character map of the wind, water, insect, vertebrate ARD model. 26 major angiosperm orders are labelled: Chlor. = Chloranthales, Cerat. = Ceratophyllales, Ranun. = Ranunculales, Faga. = Fagales, Cucu. = Cucurbitales, Solan. = Solanales, Borag. = Boraginales. Pie charts at tip labels indicate polymorphic data. The grey band indicates the period of the Cretaceous from 145 - 66 million years ago, though the timing of angiosperm evolution remains an area of active research and times displayed are indicative only.

Fifty-seven of 64 angiosperm orders were reconstructed as ancestrally insect pollinated (proportional marginal likelihoods >0.85; Figure 1). Notable exceptions include Ceratophyllales (water pollinated), Zingiberales (vertebrate pollinated), Fagales (wind pollinated) and Picramniales (wind pollinated; Figure 1). Two angiosperm clades are primarily water pollinated: Ceratophyllales and the seagrasses within Alismatales (Ruppiaceae, Cymodoceaceae, Posidoniaceae, Zosteraceae and Potamogetonaceae; Figure 1, Figure S1). Major wind pollinated clades include sections of Alismatales, Poales, Rosales, Fagales and Caryophyllales, though wind pollination occurs at many other points across the angiosperm tree (Figure 1, Figure S1). Vertebrate pollination is dispersed across the angiosperm tree, with large vertebrate pollinated clades in the Zingiberales, Bromeliaceae and at the base of the Ericales (Figure 1, Figure S1). Ancestral states inferred for all angiosperm families and orders can be seen in more detail in Supporting Information Figure S1.

Transition rates (number of transitions per million years) for the 4-state ARD model were low overall, with the highest transition rate (0.01) for reversals from vertebrate to insect pollination (Figure 2). Transition rates from insect to vertebrate pollination were an order of magnitude lower (0.0007), and rates between vertebrate pollination and wind or water pollination were near zero (<0.00001). The transition rate from insect to wind pollination (0.0007) was an order of magnitude lower than the reversal (0.001). Transition rates to water pollination were highest from wind pollination (0.002).

**Figure 2.**
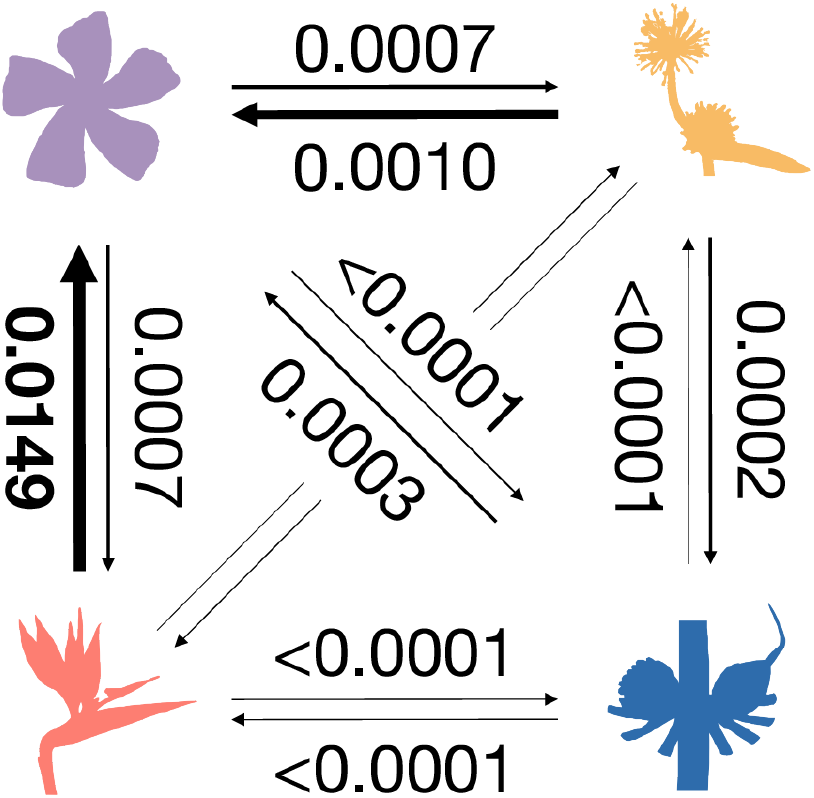
Transition matrix showing transition rates (number of transitions per million years) for the 4-state ARD model; transition rates not displayed on the diagonal between wind and vertebrate pollination are both <0.0001.

Of the 434 angiosperm families sampled, 327 (75%) were entirely animal pollinated, 37 (9%) were entirely wind pollinated, 5 (1%) were entirely water pollinated and 64 (15%) families contain a mix of taxa pollinated by wind, water and/or animals. Three hundred and eighty-nine (90%) angiosperm families contain some form of animal pollination (384 or 89% by insects) and 100 (23%) families contain some form of wind pollination including ambophily.

### 3.1 Stochastic character mapping

Stochastic character mapping found more transitions to wind pollination from animal pollination than the reverse, with a 95% Highest Posterior Density (HPD) interval of 42-50 transitions to wind and 4-12 reversals back to animal pollination (Figure 3a). In contrast, vertebrate and insect pollination had frequent transitions to and from, with 95% HPD of 39-56 transitions from insect to vertebrate pollination and 26-57 reversals from vertebrate back to insect pollination (Figure 3b). Transitions to water pollination occurred only 1-3 times, with no reversals (SI Figure S2). When averaging the total branch lengths spent in each state across all stochastic character maps, a mean 86% of angiosperm evolutionary time since the crown node is spent in insect pollination, 10% of evolutionary time in wind pollination, 4% of time in vertebrate pollination and 1% of time in water pollination.

**Figure 3.**
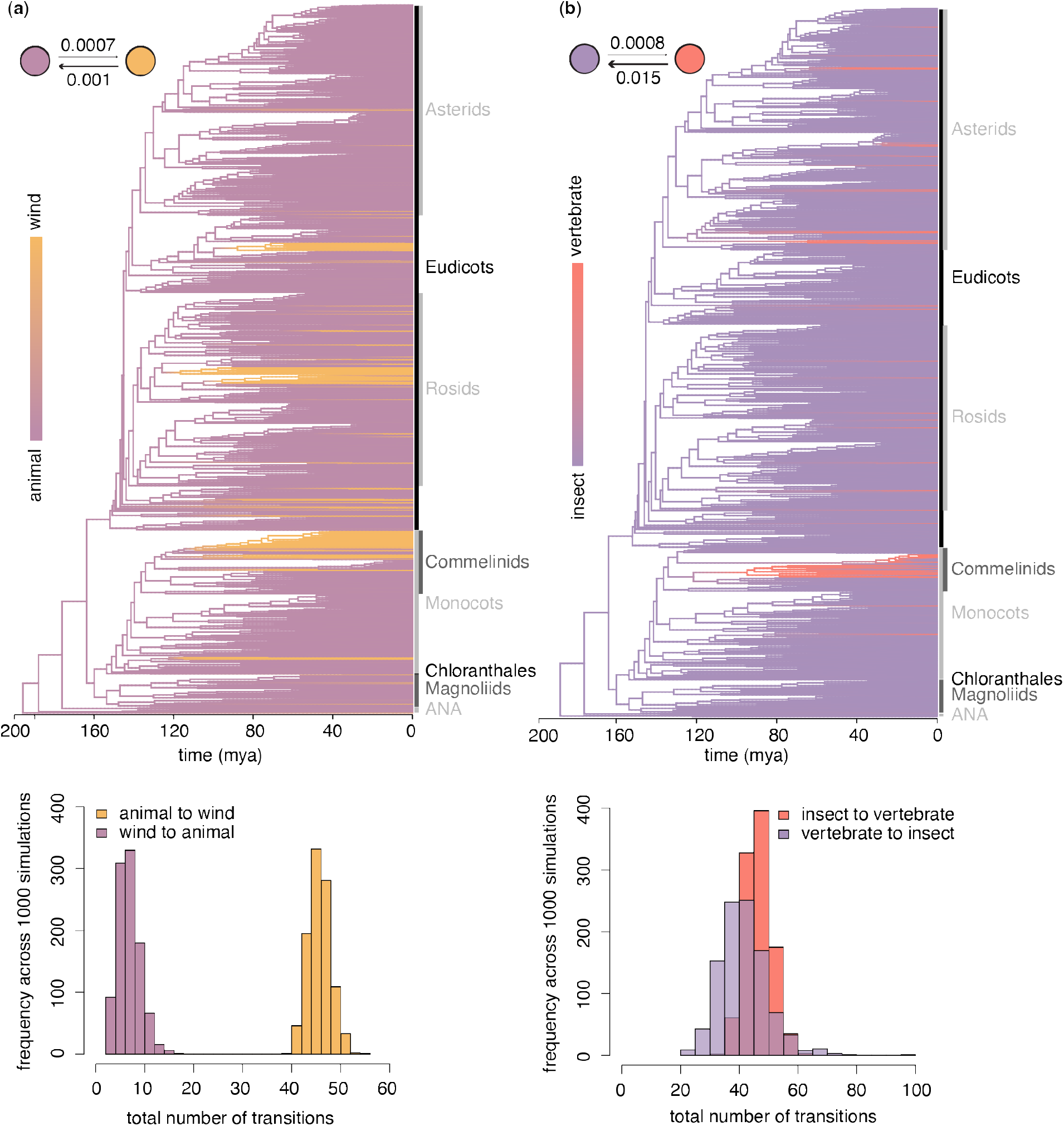
Reconstruction of transitions between wind and animal pollination (a) and, within animal pollination, between insect and vertebrate pollination (b) across the history of angiosperms. Trees are density maps from 1000 simulated stochastic histories based on ARD binary-state models; note that the tree in (b) is a subset of the tree in (a) with all wind pollinated tips removed, and water pollinated tips have been removed from both trees. Transition matrices in the top left of each panel indicate the transition rates from ancestral state reconstructions for each model. Histograms show the number of transitions between each pollination mode across all 1000 simulated stochastic histories.

Dating uncertainty around the single angiosperm tree used for above analyses had little impact on these numbers, with stochastic mapping across 1000 trees from the posterior of Ramírez-Barahona et al. (2020) returning largely congruent results (SI Table S2-S3).

The time of transitions extracted from stochastic mapping suggests that transitions to wind pollination started early in the evolution of extant angiosperms (reconstructed here as approximately 197 million years ago, (mya Figure 4a), though the age of angiosperms remains an area of active research (Sauquet *et al*., 2022)). Transitions to wind pollination start approximately 129-131 mya (95% CI), and continue steadily from then on, with a slight decrease around 80 mya followed by an acceleration towards the present day (Figure 4a). Reversals back to animal pollination are both fewer and slower than transitions to wind pollination, starting approximately 76-79 mya with most occurring in the last 40 million years (Figure 4a). Transitions between insect and vertebrate pollination track each other closely (Figure 4b). The first transition to vertebrate pollination occurs approximately 126-127 mya, with the first reversal from vertebrate back to insect pollination 103-106 mya (Figure 4b). Transitions between insect and vertebrate pollination increase sharply towards the present day (Figure 4b).

**Figure 4.**
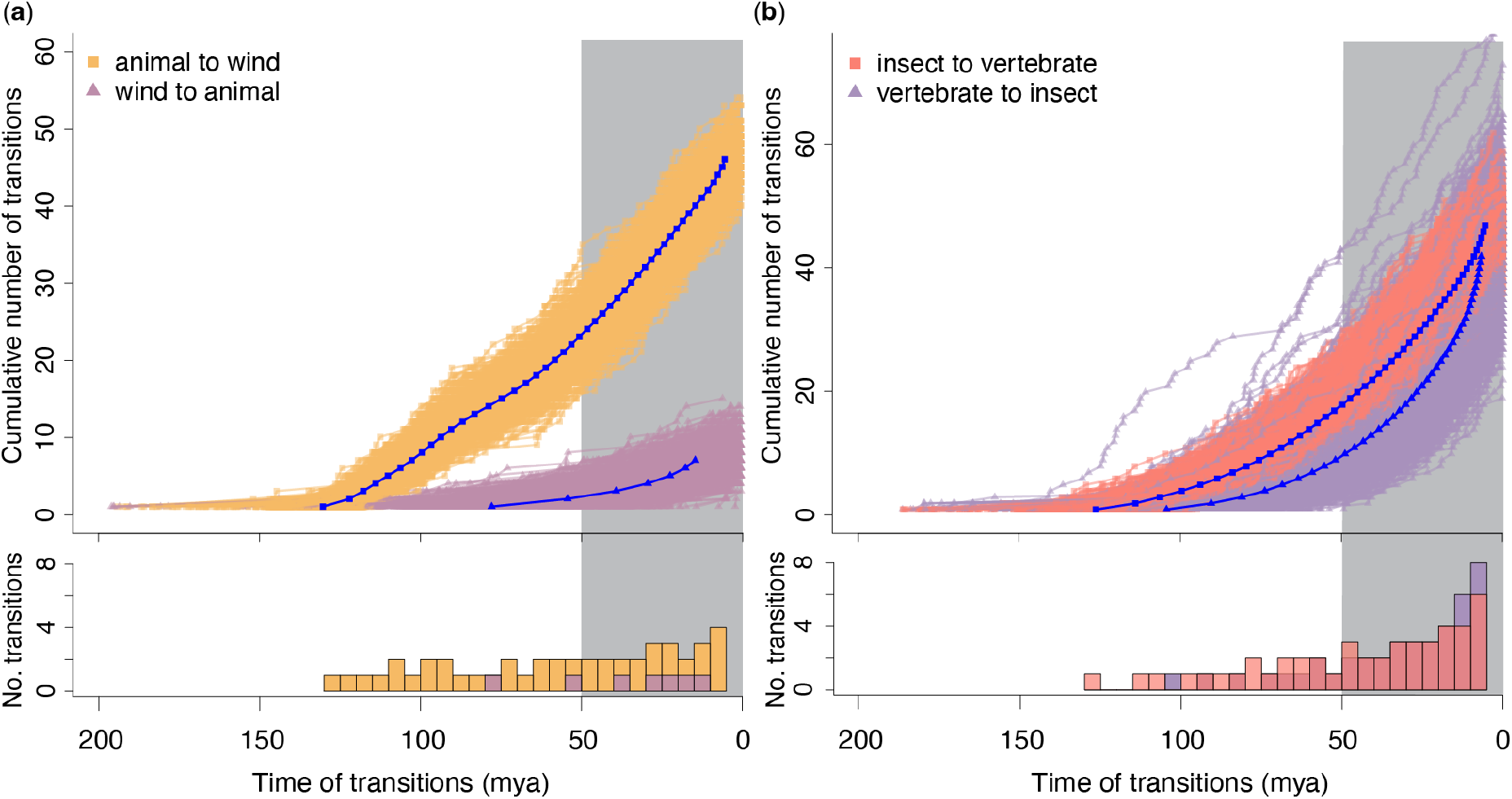
The number and timing of transitions between wind and animal pollination (a) and, within animal pollination, between insect and vertebrate pollination (b) across the history of angiosperms, based on 1000 simulated stochastic histories of ARD binary-state models. Coloured lines show individual stochastic trajectories, blue lines show the mean stochastic trajectory (timing of each transition and total number of transitions). Histograms below the main graphs show the number of transitions in each time period averaged across all simulations. The grey highlight shows the period towards the present where denser species sampling would likely detect stronger trends. The timing of angiosperm evolution remains an area of active research and times displayed are indicative only.

### 3.2 Environmental and geographic relationships

Both wind and animal pollinated species are found at a range of latitudes and across habitats which are, on average, characterised by a wide range of canopy openness values (mean LAI) (Figure 5). Species with high (~8) mean LAI were 1.2 times more likely to be animal pollinated in phylogenetic logistic regression analyses (coefficient = 0.16, confidence interval 0.16–0.27, *p* = 0.02, n = 1,022 species; Figure 5a). The probability of wind pollination increased with distance from the equator (absolute latitude), with species 2% more likely to be wind pollinated with each 1° increase in mean latitude (coefficient = −0.02, confidence interval −0.021 – −0.024, *p* = 0.03, n = 1,022 species; Figure 5b).

**Figure 5.**
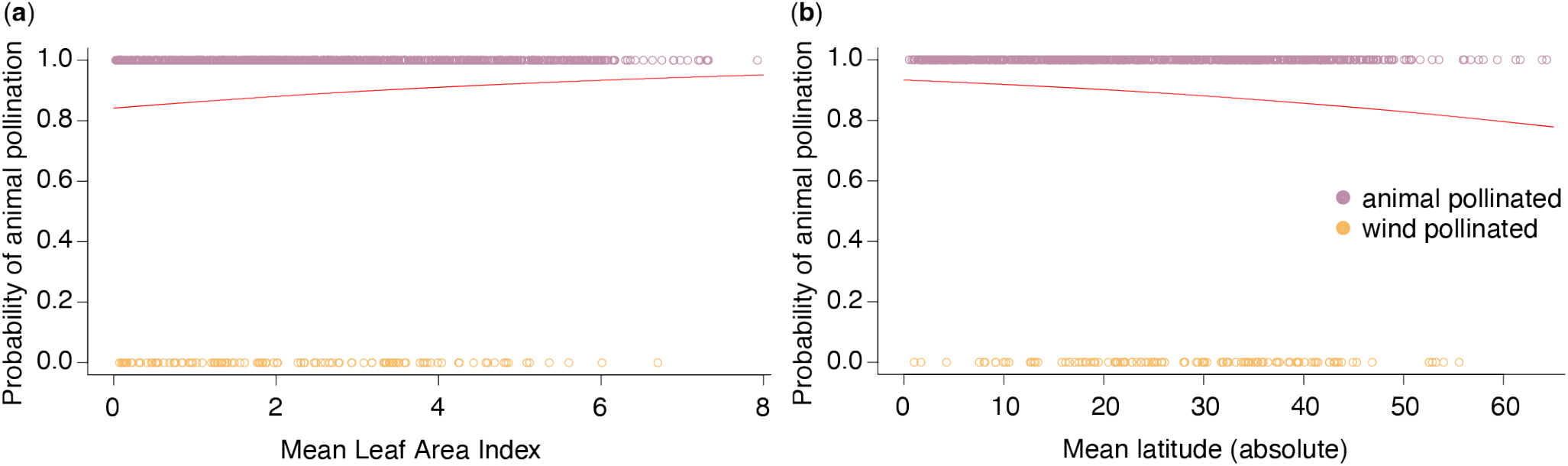
The relationship between animal/wind pollination and species mean Leaf Area Index and absolute latitude. Red lines show the coefficient of phylogenetic logistic regressions between wind/animal pollination mode and species mean Leaf Area Index (a) or species mean absolute latitude (b).

In contrast, Pagel’s models found limited evidence for correlated evolution between superbiome occupancy and wind versus animal pollination (see Supporting Information Notes S1).

## 4 Discussion

Most extant angiosperms are pollinated by insects, and our reconstructions here suggest that the most recent common ancestor of the angiosperms was also insect pollinated. This accords with previous inferences of ancestral insect pollination in the angiosperms from the fossil record (Hu *et al*., 2008; Friis *et al*., 2011; Labandeira & Currano, 2013; Asar *et al*., 2022) and reviews of pollination in early diverging angiosperm lineages (Hu *et al*., 2008; Thien *et al*., 2009; Gottsberger, 2016). Ancestral insect pollination of angiosperms is further supported by the floral syndrome of Sauquet *et al*. (2017)’s reconstructed ancestral flower: the bisexual and radially symmetric flower, with more than two whorls of perianth and stamens is consistent with a generalist insect pollination syndrome. More detail of ancestral traits such as flower size, pollen stickiness and the presence of nectaries or nectar drops would be needed to infer this with certainty, however.

Beyond just the angiosperms, evidence is accumulating that the ancestor of all seed plants may itself have been insect pollinated (Ollerton, 2017; Asar *et al*., 2022). Though many contemporary gymnosperms are wind pollinated, particularly the conifers and *Ginkgo biloba*, both extinct and extant gymnosperm lineages contain many diverse examples of insect pollination (reviewed in Labandeira & Currano, 2013; Asar *et al*., 2022). Our strong support for insect pollination at the angiosperm crown node again raises the question of how far back along the angiosperm stem insect pollination may have evolved. Definitive answers may be difficult to come by however, given the long period of angiosperm stem evolution for which we have limited fossil evidence.

Pollination by insects has clearly been a successful reproductive strategy throughout angiosperm history, with 86% of evolutionary time spent in insect pollination on average (Figure 1). Although the origins and diversification of angiosperms and insect pollinators may not be as tightly linked as previously thought (Asar *et al*., 2022), theirs is clearly a mutualism of great antiquity. This long period of interactions between pollinating insects and flowers has enabled diverse but effective pollination mutualisms to evolve (Friis *et al*., 2011), with the advantages of insect pollination for angiosperms noted by evolutionary biologists since Darwin (1876). These include the abundance of pollinating insects in a broad range of habitats, from arid deserts to Antarctic islands, though insects are increasingly abundant towards the tropics (Ollerton, 2017). Another advantage is the relatively small investment in floral rewards necessary to attract insect pollinators, especially small-bodied insects (McCallum *et al*., 2013). The dependence on floral rewards by insects such as bees or butterflies, whose entire diet is supplied by nectar and/or pollen, in turn increases insects’ flower constancy, improving the chances of pollen being transferred between compatible conspecific flowers (Grüter & Ratnieks, 2011). There are of course some drawbacks to insect pollination, especially as insects are often herbivores and eat many plant parts, not just the floral rewards provided. There have been suggestions that florivory may represent a precursor to pollination in some systems though (Xiao *et al*., 2021), and clearly the benefits of attracting insect pollinators outweigh the risks of herbivory for the majority of flowering plants.

### 4.1 Pollination transitions

We show that evolutionary shifts between insect and vertebrate pollination have been frequent throughout angiosperm history, with at least 39-56 transitions from insect to vertebrate pollination (Figure 3b). This is consistent with studies which show numerous shifts to bird, bat and other vertebrate pollination in clades such as Zingiberales and Bromeliaceae (Specht *et al*., 2012; Kessler *et al*., 2020), though there has been no angiosperm-wide assessment to our knowledge. Many of the advantages of insect pollination also apply to pollination by vertebrate animals. Vertebrate pollination can be an effective mutualism, where vertebrate dependence on floral rewards promotes flower constancy and thus efficient, targeted cross pollination (Ratto *et al*., 2018). Vertebrate pollinated flowers are often expensive for a plant to produce, needing large, robust floral parts and large quantities of pollen and nectar as floral rewards (Fleming *et al*., 2009; McCallum *et al*., 2013). In return, vertebrate pollinators can often transport pollen at greater distances than insect pollinators, thus increasing gene flow and reducing the chance of inbreeding depression for some vertebrate pollinated plants (Fleming *et al*., 2009; Wessinger, 2021). In fact, early evidence suggest that vertebrate (hummingbird) pollination is favoured in self-incompatible plants (Abrahamczyk *et al*., 2022b). It remains to be seen whether this pattern is general, such that the advantage of better outcrossing outweighs the cost of greater floral investment for vertebrate-pollinated species.

Our reconstructions suggest that reversals from vertebrate to insect pollination have been almost as frequent as the original transitions (Figure 3b). This is at odds with some studies which show strong specialisation and a lack of reversibility in vertebrate pollinated species, such as in hummingbird pollinated genera (Barrett, 2013). Other studies suggest vertebrate pollination can be more generalised, especially outside the tropics (Ratto *et al*., 2018). Indeed, our data contained numerous instances of species pollinated by both insects and vertebrates: for example, *Lapageria rosea* (Philesiaceae, Liliales) is pollinated by both hummingbirds and large bumblebees (Valdivia *et al*., 2006). Although some vertebrate pollination systems are highly specialised, others are clearly more general, and the traits separating insect from vertebrate pollination syndromes may be relatively labile. Both vertebrate and insect pollination require plants to provide floral rewards, particularly nectar, to have sticky pollen, and to attract animal pollinators with a showy perianth or similar (Faegri & van der Pijl, 1979). Floral traits separating insect from vertebrate pollination, such as flower size and colour or nectar volume, can be environmentally plastic and phylogenetically variable (e.g. McEwen & Vamosi, 2010; Parachnowitsch *et al*., 2019), suggesting that these characters may be relatively easy for plants to change. Our results question the narrative that vertebrate pollination is highly specialised and irreversible, perhaps reflecting the fact that most pollination systems are general in nature (Waser *et al*., 1996), despite the increased research attention specialisation receives.

In contrast to vertebrate and insect pollination, reversals from wind to animal pollination are rare (Figure 3a). Transitions to wind pollination require significant changes in floral traits, such as the reduction or removal of the perianth and nectaries, shift to unisexuality and dioecy, increase in the pollen:ovule ratio and changes to style morphology (Friedman & Barrett, 2009). Though none of these traits are irreversible, the tightly correlated constellation of traits may be difficult to reverse all together to return to successful cross-pollination by animals. Ambophily may play a transitional role in such instances, though ambophily has evolved more frequently from insect than wind pollinated ancestors (Abrahamczyk *et al*., 2022a). Where reversals from wind to animal pollination do occur, they typically involve a transition to generalist insect pollination (Barrett, 2013). For example, in the predominantly wind pollinated genus *Cyperus*, *C. sphaerocephalus* (Cyperaceae, Poales) has colourful bracts, pollen sticky with pollenkitt and a floral scent which attracts flies, beetles and bees, setting few seeds when insects but not wind are excluded (Wragg & Johnson, 2011). *Cyperus* flowers are mostly bisexual, and this co-occurrence of anthers and stigmas in flowers probably facilitates reversals to insect pollination in bisexual taxa (Wragg & Johnson, 2011), especially given the tendency of insects to collect pollen from wind-pollinated flowers (Saunders, 2018). In fact, though dioecy is occasionally reversible (Wang *et al*., 2021), the separation of plant sexes may be the most significant barrier to wind-animal reversals. Insects taking pollen from male flowers may never even visit female flowers, and thus are less likely to incidentally pollinate dioecious or diclinous wind-pollinated plants. Thus, reproductive barriers may contribute to the rarity of reversals from wind to animal pollination.

Despite the large number of necessary trait changes, many shifts from animal to wind pollination have occurred throughout angiosperm evolution. Our simulation of at least 42-50 transitions from animal to wind pollination accords with previous estimates of at least 65 shifts across the angiosperms (Linder, 1998), and is likely an underestimate given our subsampling approach. Major transitions to wind pollination have led to large wind pollinated clades in the Alismatales, Poales, Rosales, Fagales and Caryophyllales (Figure 1). Although it has long been suggested that wind pollination is less effective than insect pollination (e.g. Darwin, 1876), wind pollination has clearly been a successful strategy for these clades, and they have evolved diverse mechanisms to improve its efficiency (Friedman & Barrett, 2009). What drove these shifts to wind pollination remains unclear, though our exploration of environmental correlates provides some clues. Wind pollination is believed to evolve when animal pollination is limited or unreliable and the abiotic environment is conducive to wind flow (Culley *et al*., 2002; Friedman & Barrett, 2009). Significant habitat and climatic changes through angiosperm history may have opened habitats and disrupted access to pollinators, not least the Cretaceous-Paleogene extinction event (Asar *et al*., 2022). Indeed, we find wind pollination is more probable in open habitats with low Leaf Area Index as well as locations further from the equator (Figure 5), both places with higher wind flow and lower pollinator activity (Rech *et al*., 2016; Ollerton, 2017). We also find tentative evidence for correlated evolution between shifts to wind pollination and arid biomes, though deeper sampling of wind pollinated arid taxa is needed to confirm this relationship (SI). Whether wind pollination preceded and enabled angiosperm shifts to arid, open, extra-tropical habitats or evolved in response to these shifts remains an open question, though the numerous animal pollinated taxa in similar habitats suggest that wind pollination is not a precondition for such shifts.

### 4.2 Transition timing

The age of angiosperms remains highly controversial, and the evolutionary times presented here are one amongst many possible scenarios which diverge by up to 130 million years (Sauquet *et al*., 2022). Thus the timing of pollination shifts through angiosperm history remain highly uncertain, though fossil flowers, pollen and pollinators can provide some evidence (Friis *et al*., 2011). Fossil evidence for insect-angiosperm pollination extends as far back as the mid-Cretaceous (99 mya, Bao *et al*., 2019). Our models suggest the first shifts to wind pollination leading to major wind pollinated lineages occurred between 130-80 mya (Figure 3a, 4a). Fossil pollen supports angiosperm wind pollination since at least the Cenomanian (~100-95 mya, Hu *et al*., 2008), and early shifts to wind pollination could have been spurred by major climatic disruptions to pollinators throughout the Cretaceous (e.g. Linnert *et al*., 2014). Our reconstruction suggests the first shift to vertebrate pollination occurred approximately 126-127 mya, significantly earlier than fossil evidence for specialised bird or bat interactions with flowers. The earliest evidence for flower visiting behaviour in a bird dates back to the Eocene 48 mya (Mayr & Wilde, 2014), and the oldest nectar feeding bat family (Pteropodidae) originated approximately 56 mya (Fleming *et al*., 2009). Specialised vertebrate pollinators were likely preceded by generalist, opportunistic flower visitors, but how far back generalist vertebrate pollination extends may be difficult to determine, especially given larger, vertebrate pollinated flowers are less likely to be preserved in the fossil record.

### 4.3 Limitations and future directions

We provide here an overview of major evolutionary patterns of pollination across the angiosperms. Trait-independent subsampling of the phylogeny allows us to characterise patterns independent of possible biases in the literature (e.g. Adamo *et al*., 2021), and any patterns observed, such as the low number of transitions to water pollination, are likely a subset of the patterns that would emerge from denser sampling. We envision future work will add detail to our broad overview, especially as the phylogeny of angiosperms becomes ever more resolved. Moreover, our understanding of angiosperm pollination depends on the fundamental work of pollination ecologists to document complex and sometimes cryptic pollination systems. Thorough pollination studies that explicitly test for cryptic pollination mechanisms including selfing and ambophily are still needed in many angiosperm families (Ollerton, 2017; Abrahamczyk *et al*., 2022a). Further pollination studies will enable us to more fully understand the diversity and macroevolutionary dynamics of pollination systems globally.

## CONCLUSION

The mutualistic relationship between angiosperms and insect pollinators is likely ancestral and has been maintained for approximately 86% of angiosperm evolutionary history. Moreover, with at least 89% of contemporary angiosperm families insect pollinated, the relationship between plants and insect pollinators is clearly important to plant reproduction and persistence today. How pollination will continue in the Anthropocene remains to be seen.

## Supporting information

Supplementary data references

## Data availability

All data and R code needed to re-create analyses are available on GitHub at https://github.com/rubysaltbush/pollination-macroevolution and at https://doi.org/10.5281/zenodo.7592528.

## Acknowledgements

We thank Santiago Ramírez-Barahona for providing superbiome occupancy data and the posterior trees for stochastic mapping. We are grateful to A. López-Martínez, J. Baczyński and J. Herting for help with macroevolutionary and data visualisation methods, and to the Sauquet lab for many stimulating discussions about macroevolution, flowers and pollination. R.E.S. was supported by funding through the Australian Government’s Research Training Program.

## Competing interests

The authors declare no conflict of interest.

## Author contributions

R.E.S., H.S. and R.V.G. conceived the project; R.E.S. and L.D. assembled the pollination data; W.C. assisted with geographic data and analyses. R.E.S. analysed the data; R.E.S. led the writing with assistance and review from all other authors.

**The following Supporting Information is available for this article:**

**Fig. S1** Tall phylogeny showing the macroevolution of pollination modes across angiosperms.

**Fig. S2** Transitions to wind, water, vertebrate and insect pollination across 1000 simulations.

**Fig. S3** Species latitude versus Leaf Area Index, coloured by superbiome occupancy.

**Table S1** Ancestral State Reconstruction models for different pollination mode categorisations.

**Table S2** Number of transitions between pollination states for main tree versus posterior trees.

**Table S3** Proportion tree time in pollination states for main tree versus posterior trees.

**Table S4** Number of species sampled by pollination mode and superbiome.

**Table S5** Model results for correlated evolution between pollination and superbiomes.

**Notes S1** Testing for correlated evolution between pollination mode and superbiome occupancy.

## Notes

https://doi.org/10.5281/zenodo.7592528

### Competing Interest Statement

The authors have declared no competing interest.

