## Supporting Information for "Insects pollinated flowering plants for most of angiosperm evolutionary history"

The following Supporting Information is available for this article:

**Fig. S1** Tall phylogeny showing the macroevolution of pollination modes across angiosperms.

**Fig. S2** Transitions to wind, water, vertebrate and insect pollination across 1000 simulations.

**Fig. S3** Species latitude versus Leaf Area Index, coloured by superbiome occupancy.

**Table S1** Ancestral State Reconstruction models for different pollination mode categorisations.

**Table S2** Number of transitions between pollination states for main tree versus posterior trees.

**Table S3** Proportion tree time in pollination states for main tree versus posterior trees.

**Table S4** Number of species sampled by pollination mode and superbiome.

**Table S5** Model results for correlated evolution between pollination and superbiomes.

**Notes S1** Testing for correlated evolution between pollination mode and superbiome occupancy.

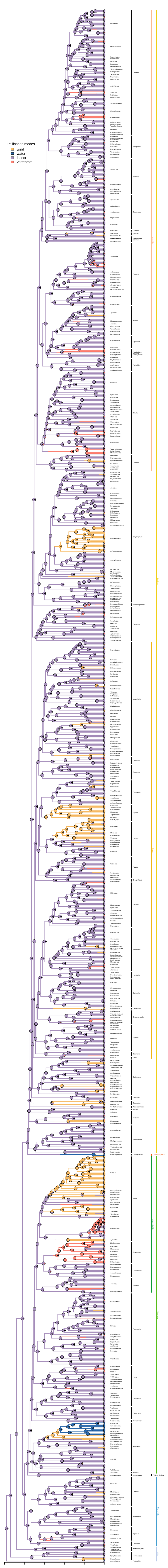

Pollination modes
wind
water
insect
vertebrate
Araceae
Tofieldiaceae
Acoraceae
Acorales
Chloranthaceae
Chloranthales
Chloranthales
Lauraceae
Monimiaceae
Hernandiaceae
Atherospermataceae
Gomortegaceae
Siparunaceae
Calycanthaceae
Laurales
Annonaceae
Eupomatiaceae
Magnoliaceae
Degeneriaceae
Himantandraceae
Myristicaceae
Magnoliales
Magnoliids
Piperaceae
Saururaceae
Aristolochiaceae
Piperales
Winteraceae
Canellaceae
Canellales
Schisandraceae
Trimeniaceae
Austrobaileyaceae
Austrobaileyales
Cabombaceae
Nymphaeaceae
Hydatellaceae
Nymphaeales
Amborellaceae
Amborellales
ANA
200
160
120
80
40
0
time (mya)

**Fig. S1** The macroevolution of pollination modes across angiosperms, as for Figure 1 in the main text but in a long format to show more detail. The proportional marginal likelihood of pollination mode at all ancestral nodes is shown from the 4-state ARD model. Colour along tree branches indicates pollination mode from one randomly selected 4-state ARD model stochastic character map. All angiosperm families, orders and major groups are labelled. Pie charts at tip labels indicate polymorphic data.

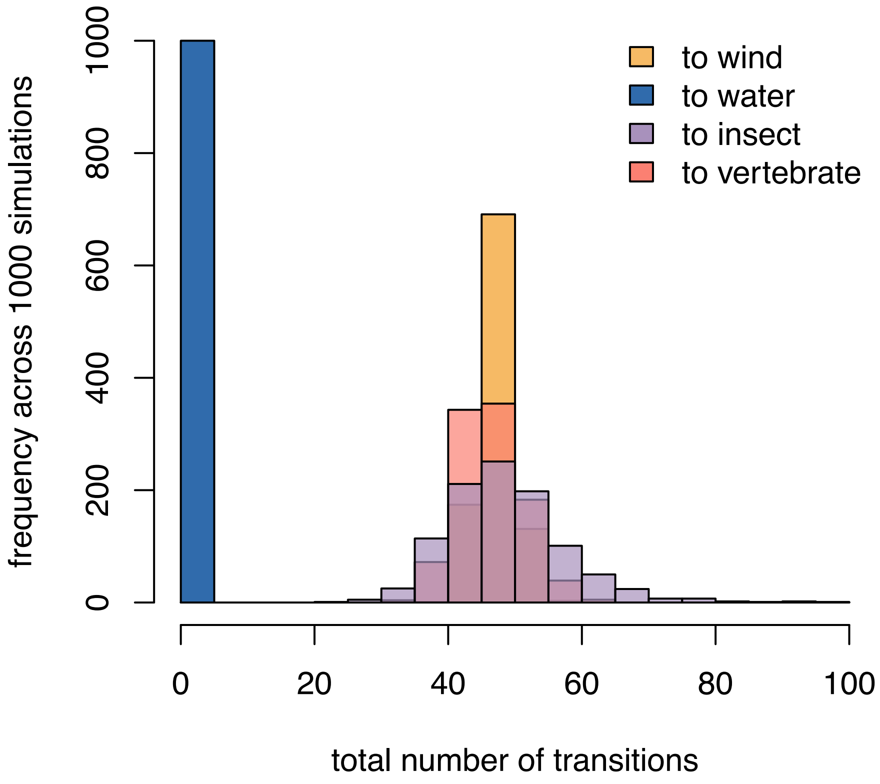


**Fig. S2** Total number of transitions to wind, water, vertebrate and insect pollination across 1000 stochastic simulations based on a wind-water-vertebrate-insect 4-state ARD model.

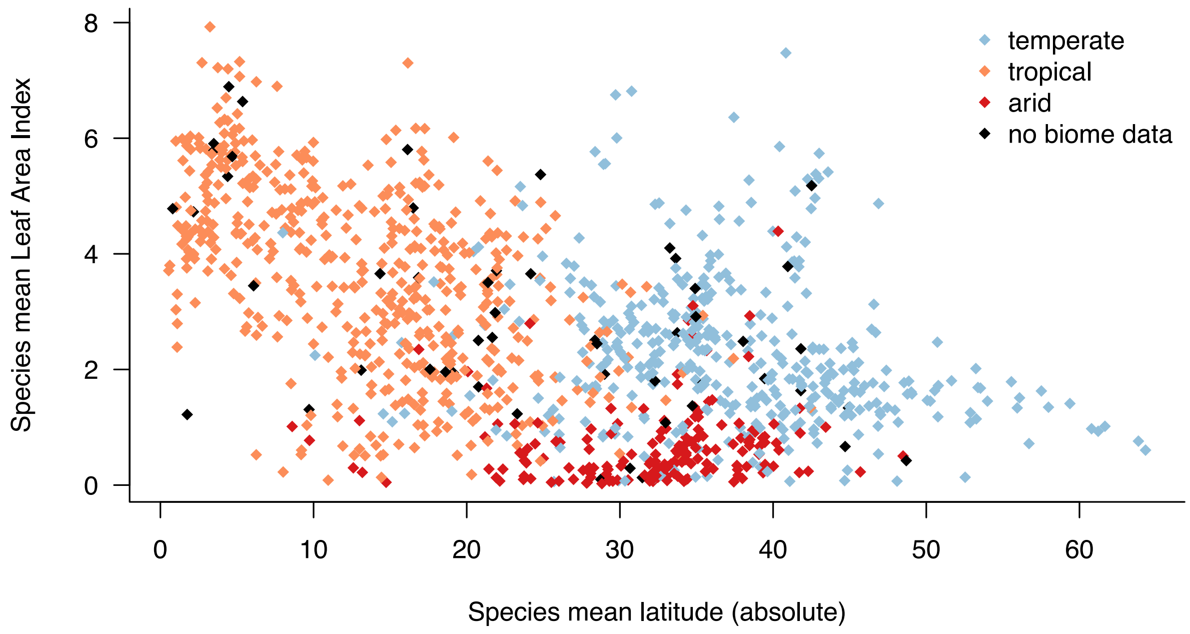


**Fig. S3** Species mean absolute latitude versus species mean Leaf Area Index, coloured by superbiome occupancy as per Ramírez-Barahona et al., 2020.

**Table S1** Ancestral State Reconstruction models fitted for different categorisations of pollination modes. Models with a+b states allow possibility of ancestral polymorphisms. The best models, chosen by lowest AICc, are marked in bold. The ancestral angiosperm state gives the most probable pollination mode for that model at the root of the angiosperm phylogeny with its proportional marginal likelihood /1. ER = Equal Rates, ARD = All Rates Different.

| **Pollination modes** | **Rate parameters** | **lnLikelihood** | **No. of parameters** | **AIC$_c$** | **Ancestral angiosperm state (proportional marginal likelihood)** |
|---|---|---|---|---|---|
| Abiotic, animal | ER | -208.9 | 1 | 419.8 | Animal (0.993) |
| | ARD | -208.8 | 2 | 421.7 | Animal (0.992) |
| Abiotic, animal, abiotic+animal | ER | -450.4 | 1 | 902.8 | Animal (0.94) |
| | **ARD** | **-436.8** | **6** | **885.6** | **Animal (0.995)** |
| Wind, water, animal | ER | -250.0 | 1 | 502 | Animal (0.996) |
| | **ARD** | **-221.0** | **6** | **454.2** | **Animal (0.993)** |
| Wind, water, animal, wind+water, wind+animal, water+animal | ER | -562.4 | 1 | 1126.8 | Animal (0.953) |
| | **ARD** | **-461.8** | **30** | **985.1** | **Animal (0.953)** |
| Abiotic, insect, vertebrate | ER | -387.5 | 1 | 777 | Insect (0.992) |
| | **ARD** | **-367.3** | **6** | **746.7** | **Insect (0.979)** |
| Abiotic, insect, vertebrate, abiotic+insect, abiotic+vertebrate, insect+vertebrate | ER | -938.6 | 1 | 1879.2 | Insect (0.937) |
| | **ARD** | **-831.1** | **30** | **1723.9** | **Insect (0.952)** |
| Wind, water, insect, vertebrate | ER | -429.5 | 1 | 861 | Insect (0.995) |
| | **ARD** | **-379.5** | **12** | **783.4** | **Insect (0.981)** |

**Table S2** 95% Highest Posterior Density (HPD) intervals for the number of transitions between different states when stochastically simulated 1000 times on the main 'RC-complete' tree and 100 times each across 1000 posterior angiosperm trees from Ramírez-Barahona et al. (2020) to consider the impact of dating uncertainty on results.

| **Model** | **Transition** | **95% HPD number of transitions** | |
|---|---|---|---|
| | | RC-complete tree | Posterior trees |
| Wind-animal ARD | Animal to wind | 42-50 | 41-51 |
| Wind-animal ARD | Wind to animal | 4-12 | 3-13 |
| Insect-vertebrate ARD | Insect to vertebrate | 39-56 | 40-61 |
| Insect-vertebrate ARD | Vertebrate to insect | 26-57 | 28-66 |
| 4-state ARD | Wind to water | 1-2 | 0-2 |
| 4-state ARD | Wind to insect | 3-11 | 3-13 |
| 4-state ARD | Wind to vertebrate | 0 | 0-1 |
| 4-state ARD | Water to wind | 0 | 0-1 |
| 4-state ARD | Water to insect | 0 | 0 |
| 4-state ARD | Water to vertebrate | 0 | 0 |
| 4-state ARD | Insect to wind | 43-52 | 12-52 |
| 4-state ARD | Insect to water | 0-1 | 0-1 |
| 4-state ARD | Insect to vertebrate | 38-56 | 37-75 |
| 4-state ARD | Vertebrate to wind | 0 | 0-33 |
| 4-state ARD | Vertebrate to water | 0 | 0-1 |
| 4-state ARD | Vertebrate to insect | 26-59 | 25-209 |

**Table S3** Mean proportion of total time (branch lengths from crown node) spent in each state for stochastic character mapping of 1000 simulations on the main 'RC-complete' tree and 100 simulations across 1000 posterior angiosperm trees from Ramírez-Barahona et al. (2020) to consider the impact of dating uncertainty in the angiosperm tree on results.

| **Model** | **State** | **Mean proportion total time** | |
|---|---|---|---|
| | | RC-complete tree | Posterior trees |
| Wind-animal ARD | Wind | 0.10 | 0.10 |
| Wind-animal ARD | Animal | 0.90 | 0.90 |
| Insect-vertebrate ARD | Insect | 0.96 | 0.96 |
| Insect-vertebrate ARD | Vertebrate | 0.04 | 0.04 |
| 4-state ARD | Wind | 0.10 | 0.10 |
| 4-state ARD | Water | 0.01 | 0.01 |
| 4-state ARD | Insect | 0.86 | 0.85 |
| 4-state ARD | Vertebrate | 0.04 | 0.05 |

**Table S4** Number of species sampled in our data by pollination mode (wind vs. animal) and superbiome occupancy (tropical, temperate or arid).

| Pollination mode | Superbiome occupancy | Number of species |
|---|---|---|
| Animal | Tropical | 440 |
| Animal | Temperate | 322 |
| Animal | Arid | 143 |
| Animal | Missing | 58 |
| Wind | Tropical | 41 |
| Wind | Temperate | 43 |
| Wind | Arid | 24 |
| Wind | Missing | 9 |
| Wind & animal | Tropical | 23 |
| Wind & animal | Temperate | 29 |
| Wind & animal | Arid | 11 |
| Wind & animal | Missing | 3 |
| Missing | Tropical | 21 |
| Missing | Temperate | 8 |
| Missing | Arid | 2 |
| Missing | Missing | 10 |

**Table S5** Model parameters and results for Pagel's (1994) models of correlated evolution between wind/animal pollination mode and different categories of superbiome occupancy across the angiosperms. All models run with equal weight root priors and ten random restarts. The best models, chosen by AICc, are in bold.

| Superbiome occupancy model | Evolutionary model | lnLikelihood | No. of parameters | $AIC_c$ | $\Delta AIC_c$ |
|---|---|---|---|---|---|
| **Tropical, temperate, arid** | **Uncorrelated** | **-1266** | **18** | **2549** | **0** |
| Tropical, temperate, arid | Correlated | -1258 | 8 | 2553 | 4 |
| Tropical, non-tropical | Correlated | -910 | 8 | 1836 | - |
| Tropical, non-tropical | Uncorrelated | -914 | 4 | 1836 | - |
| Temperate, non-temperate | Correlated | -897 | 8 | 1811 | - |
| Temperate, non-temperate | Uncorrelated | -901 | 4 | 1810 | - |
| **Arid, non-arid** | **Correlated** | **-683** | **8** | **1383** | **0** |
| Arid, non-arid | Uncorrelated | -691 | 4 | 1389 | 6 |

**Notes S1** Testing for correlated evolution between pollination mode and superbiome occupancy.

While exploring the environmental relationships of wind pollination in our data, we tested whether the evolution of wind pollination is correlated with evolutionary shifts of angiosperms between different major biomes using superbiome occupancy data from Ramírez-Barahona *et al.* (2020). Superbiome data assign each species in our study as tropical, temperate or arid based on the occurrence of at least 50% of cleaned GBIF records in that superbiome (Figure S5). Species superbiomes approximately correspond with species mean LAI and absolute latitude as per our other measures of environmental relationships (Figure S5, Figure 5 main text).

We used corHMM to implement Pagel's (1994) maximum likelihood method of testing for correlated evolution, using all three superbiomes together as well as three binary categorisations (tropical/non-tropical, temperate/non-temperate and arid/non-arid) to reduce the number of model parameters. Fourteen water pollinated taxa were excluded from evolutionary correlations, giving a final sample size n = 1187 (Table S4). To confirm these relationships we also ran phylogenetic logistic regressions for each superbiome category versus wind/animal pollination, fitted in phylolm version 2.6.2 (Tung Ho & Ané, 2014) following the Ives and Garland method (Ives & Garland, 2010). Uncertainty of parameter estimates was estimated by 100 parametric bootstrap samples. Taxa with missing or polymorphic data as well as water pollinated taxa were excluded from phylogenetic logistic regressions, giving a final sample size of n = 1013.

Pagel's models do not support correlated evolution between tropical, temperate and arid superbiome occupancy and wind/animal pollination (**$\Delta AIC_c$** = 4, Table S5). Support was found for correlated evolution between arid and non-arid superbiome occupancy and wind/animal pollination (**$\Delta AIC_c$** = 6, Table S5), though the estimated transition rates for this correlated model were much higher than would be expected (27-100 when usually <1), suggesting that this model was affected by potential estimation artefacts linked with rare states, with only 24 wind-pollinated arid superbiome taxa in our data (Tables S4-S5). Phylogenetic logistic regressions found no relationship between wind pollination and superbiome occupancy ($p > 0.1$).

These mixed results do not provide strong support for correlated evolution between pollination mode and superbiome occupancy. The arid/non-arid models were likely affected by undersampling in our data, and deeper sampling with a phylogeny that includes more wind-pollinated arid taxa would be needed to confirm whether the support found for correlated evolution here is genuine. It is also worth noting that the superbiome categories used here are necessarily broad, simplifying the vast range of continuous environmental variation experienced by each taxon in our data into only three general categories. The tropical superbiome, for example, covers a broad range of Leaf Area Indices (Figure S5) as it includes both closed tropical rainforest and open tropical savannah biomes. We hope that future work can clarify the relationship between evolutionary shifts in pollination mode and environment, either with deeper sampling or a more targeted approach.
